# AWET – Arthropod Weight Estimation Tool

**DOI:** 10.64898/2026.09.03.749054

**Authors:** Mathieu Cretton, Ruedi Boesch, Janine Bolliger, Kurt Bollmann, Roman Flury, Malte Jochum, Nicola van Koppenhagen, Thomas Zimmermann, Martin K. Obrist

**Affiliations:** WSL, Swiss Federal Research Institute, Zürcherstrasse 111, CH-8903 Birmensdorf, Switzerland; Department of Global Change Ecology, Biocenter, University of Würzburg, Emil-Hilb-Weg 22, 97074 Würzburg, Germany; Department of Biology, ETH Zürich, Wolfgang-Pauli-Strasse 27, CH-8093 Zürich, Switzerland; Department of Biosystems Science and Engineering, ETH Zürich Schanzenstrasse 44, CH-4056 Basel, Switzerland

**Keywords:** allometric relationships, arthropod biomass, automated image analysis, body weight, body size, ImageJ libraries

## Abstract

Arthropods drive essential ecosystem processes such as pollination, decomposition, and nutrient cycling and are widely used as indicators of ecosystem condition and function. Among various arthropod-derived metrics, body weight is a key variable in functional ecology and frequently assessed as dry body weight. However, drying arthropod specimens limits the samples’ future potential for research, as it prevents further processing such as trait measurements or species identification.

Here, we present AWET (Arthropod Weight Estimation Tool), an open-source application for automated estimation of individual fresh body weight and extraction of morphometric measurements from standardized images of pre-sorted arthropod samples. AWET combines automated image analysis with taxon-specific allometric regression models to estimate fresh body weight while simultaneously quantifying body length, width, area, and specimen abundance. The software operates with standard imaging equipment, requires no machine-learning-based classification or segmentation, and allows users to define taxonomic groupings according to their objectives. By preserving specimens for downstream analyses while processing large numbers of individuals within milliseconds, AWET provides an efficient, non-destructive, and cost-effective workflow for high-throughput arthropod phenotyping. The software is a practical and expandable tool for biodiversity monitoring projects investigating changes in arthropod biomass, abundance and individual morphometric measures.

## 1. Introduction

The ongoing decline of arthropods has obtained global attention in the last decade (Hallmann et al., 2017), particularly with respect to losses in abundance, richness, and biomass (Emer et al., 2019; Hallmann et al., 2021). Arthropods are vital in providing ecologically relevant functions and services that are crucial in maintaining ecosystem processes and health (Noriega et al., 2018). Arthropods are also key for food webs as they constitute an important food source for many organisms of higher trophic levels and serve as indicators of food web structure (Brose et al., 2019). However, arthropods can also cause ecosystem disservices (Vansynghel et al., 2022), including crop damage (Trębicki et al., 2017) or vector borne diseases (Shaw & Catteruccia, 2019). Owing to their diverse and complex role in ecosystems, arthropods are widely used as indicator for monitoring ecosystem processes (Chowdhury et al., 2023).

Beyond species richness and abundance, body size and body weight represent fundamental functional traits that strongly influence metabolism, trophic interactions, and population dynamics (Kalinkat et al., 2015). Changes in habitat quality, land use, and climate are expected to alter not only the abundance of arthropods but also their size distributions and biomass, making these parameters increasingly important for ecological monitoring and biodiversity assessment (Fürst et al., 2023). Reliable estimation of individual body weight and morphometric traits is therefore becoming an essential component of studies investigating ecosystem functioning, and community responses to environmental change.

Despite this growing demand, measuring arthropod biomass remains labor-intensive and often requires destructive procedures. This limits the number of specimens that can be processed while preventing subsequent taxonomic or molecular analyses. Recent advances in digital imaging and automated image analysis have considerably improved the efficiency of arthropod phenotyping. Several software tools have been developed to extract morphometric traits or estimate biomass from digital images (Araújo Foerster et al., 2024; Kendall et al., 2019; Penell et al., 2018; Shimazaki et al., 2022), and dedicated imaging platforms increasingly integrate robotics and machine learning to automate specimen processing (Ärje et al., 2020; S. Schneider et al., 2022; Shirali et al., 2026; Wührl et al., 2022). However, most existing approaches are limited to specific taxonomic groups, require specialized imaging systems, trained machine-learning models, or focus on morphometric analysis rather than rapid and high-throughput estimation of fresh body weight. Consequently, a practical and broadly applicable workflow for estimating individual fresh body weight across diverse arthropod taxa using standard laboratory equipment remains lacking.

To address this gap, we developed AWET (Arthropod Weight Estimation Tool), an open-source application that combines automated image analysis with allometric regression models to estimate individual fresh body weight and extract morphometric measurements from pre-sorted arthropod specimens. The software operates with standardized images acquired using standard laboratory imaging equipment. It requires no machine-learning-based classification or segmentation and preserves specimens for subsequent taxonomic or molecular investigations. In addition to estimating abundance and fresh body weight, AWET quantifies body length, width, circumference, and area, thereby providing standardized, individual-level trait data suitable for ecological monitoring and biodiversity research. By combining rapid processing, flexible taxonomic grouping, and an accessible workflow, AWET aims to facilitate high-throughput quantification of arthropod biomass and morphometrics across a wide range of ecological applications.

## 2. Material and Methods

### 2.1. Allometric regression models

Using R (R Core Team, 2024), we developed allometric regression models based on the temperate arthropod dataset (N = 3’060; juveniles and larvae were excluded) and allometric relationships from Sohlström et al. (2018). These regression models allow the estimation of arthropod biomass (Fig. 1A–B) for 25 taxonomic groups (Appendix A1, Table A1.1). Some arthropod groups represent single families, while others combine multiple families based on morphological similarity. However, the latter can be further refined using the provided R script, which allows users to customize taxonomic groupings by adding, removing, or reallocating families among groups according to their specific analytical needs. Unlike Sohlström et al. (2018) who predicted body weight from body length and maximum body width, our approach specifically tested models using body length, maximal body width, and body area. Accordingly, represents the fresh body weight of a specimen for all *n* specimen in a group, then the statistical model is defined as

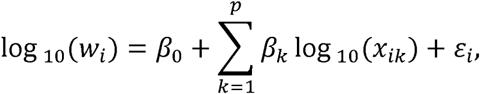

where *β*_0_ denotes the intercept, *β_k_* are regression coefficients for predictor *k* of total *p* predictors,*x_ik_* is the value of predictor *k* for specimen *i*, and ε_i_ is the residual error term. The residual errors were assumed to be independent and identically distributed

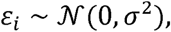

with mean zero and constant variance σ^2^. Model parameters were estimated using ordinary least squares in R. Prior to analysis, all variables, including the variable of interest, were log_10_-transformed, to meet the assumption of homoscedasticity. Body length and width measurements were recorded in millimeters [mm], body area in square millimeters [mm²], and fresh weight in milligrams [mg]. Multicollinearity among predictors was evaluated using variance inflation factors (VIF). Models with VIF values larger than five were excluded from further analysis. This procedure reduced instability in coefficient estimates, limited variance inflation, and enhanced the identifiability of individual predictor effects within the model (James et al., 2013). For each taxonomic group, all admissible combinations of predictors were fitted as linear regression models. Observations with missing values in the variables required for a given model were excluded from that model. Model performance was evaluated using the Akaike Information Criterion (AIC), Bayesian Information Criterion (BIC), coefficient of determination (*R*^2^), adjusted *R*^2^ and the root mean squared error (RMSE).

**Figure 1.**
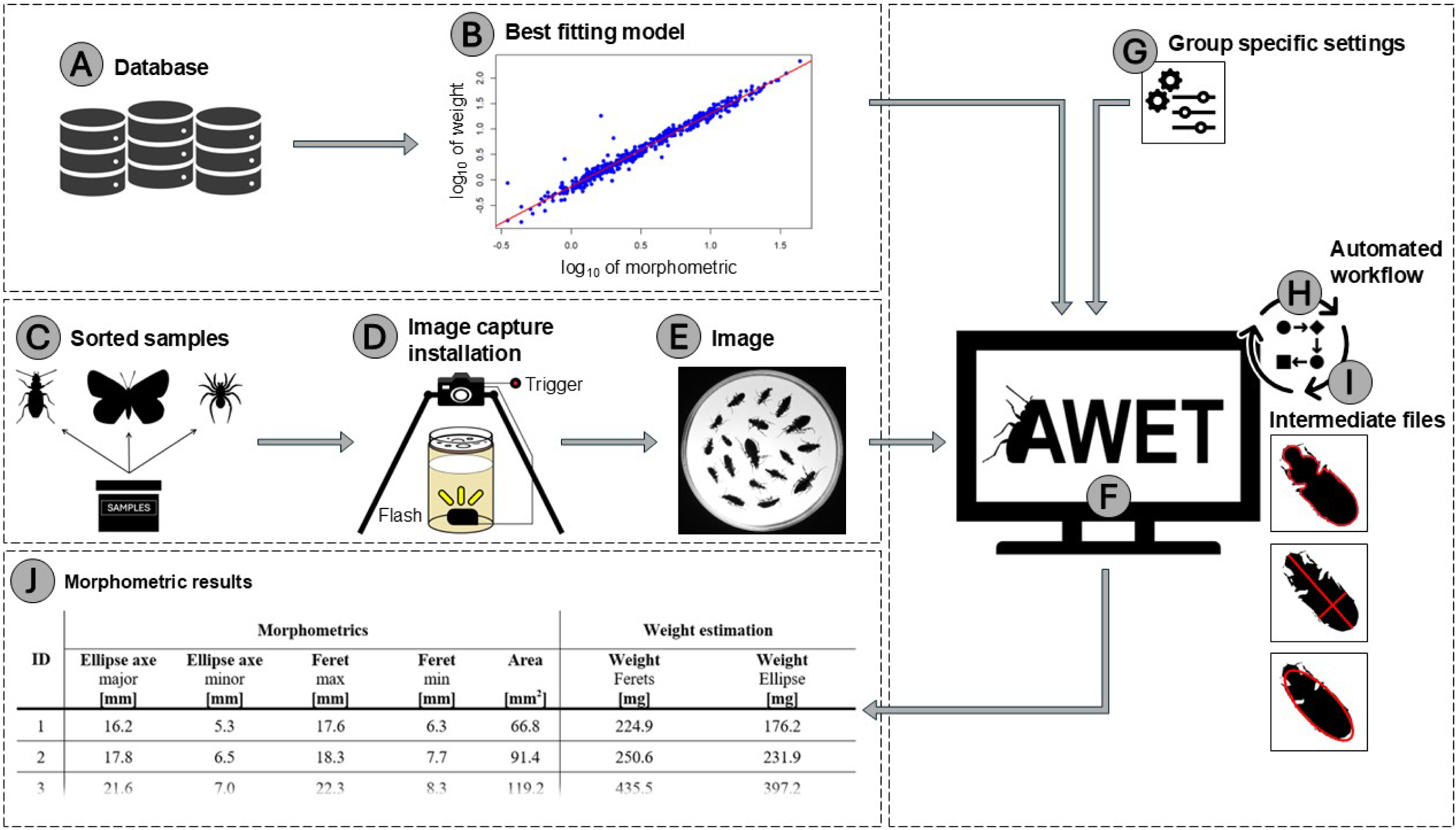
Workflow of the analytical pipeline. (A) With the dataset of temperate arthropods of Sohlström et al. (2018) we developed taxonomic arthropod group-specific allometric regression models (B). In parallel, field samples, sorted into taxonomic arthropod groups (C), were photographed (D) in a standardized way (E). Both the corresponding regression models (B) and images (E) were then processed in the AWET application (F) using group-specific parameter settings (G). The automated image-processing workflow (H) generates intermediate images and region-of-interest files, from which morphometric traits (red lines) are extracted (I). Finally, these measurements are used to estimate individual body weight of each specimen within an arthropod group. Results are exported in a CSV file (J).

Model fits were evaluated using residual diagnostics and plots of observed versus fitted values. The regression models (Appendix A2, Table A2.1) were then implemented in AWET (Fig. 1F), a Java-based application built on ImageJ libraries (C. A. Schneider et al., 2012) for the later described weight estimation.

The list of linear regression models is made available to allow the choice of the best models for specific study cases. Although the models are ranked according to their BIC and model fit, the final choice of model should also consider ecological relevance. The top-ranked model should generally be tested first, but the resulting body weight or biomass estimates should be checked to ensure that they are biologically realistic for the taxonomic group being analyzed. If the highest-ranked model produces unrealistic values, a lower-ranked model may be more appropriate. In some cases, different models may be suitable for the estimations (Appendix A3).

### 2.2. Standardized images

Standardized images of pre-sorted arthropod taxonomic samples are taken (Fig. 1C-E). The setting of the image capture installation is described in Appendix A5. Before taking the images, the specimens of a pre-sorted sample of arthropods are regularly distributed across a Petri dish. All specimens need to be clearly separated from each other for individual distinction (Figs 1E & 2A). At this stage, scaling is essential to ensure accurate morphometric measurements. The scaling procedure is typically performed by first acquiring an image containing a calibration scale (e.g. metric ruler) under the same camera settings and distance used for the specimens. The known distance on the ruler is then measured in pixels using image analysis software such as ImageJ and a conversion factor (pixels per millimeter) is calculated and applied to all subsequent measurements. This approach ensures that all derived size estimates are standardized and comparable across images and datasets.

### 2.3. Image processing

The standardized images are uploaded onto AWET to start the weight estimation. To account for differences in arthropod morphology and coloration, group specific settings (Fig. 1G; Appendix A4) are applied. The automated image-processing workflow (Fig. 1H-I) then generates four intermediate images corresponding to successive steps (Fig. 2B-E). The original image (Fig. 2A) is first converted to grayscale (Fig. 2B) to standardize pixel intensity values. Second, an optional flat-field correction is then applied to correct spatial heterogeneity in illumination and sensor response (Fig. 2C). Third, image segmentation is subsequently performed via thresholding to produce a binary image separating foreground objects from the background (Fig. 2D). Finally, small artefacts and noise are removed, resulting in a filtered binary image without outlier pixels (Fig. 2E).

**Figure 2.**
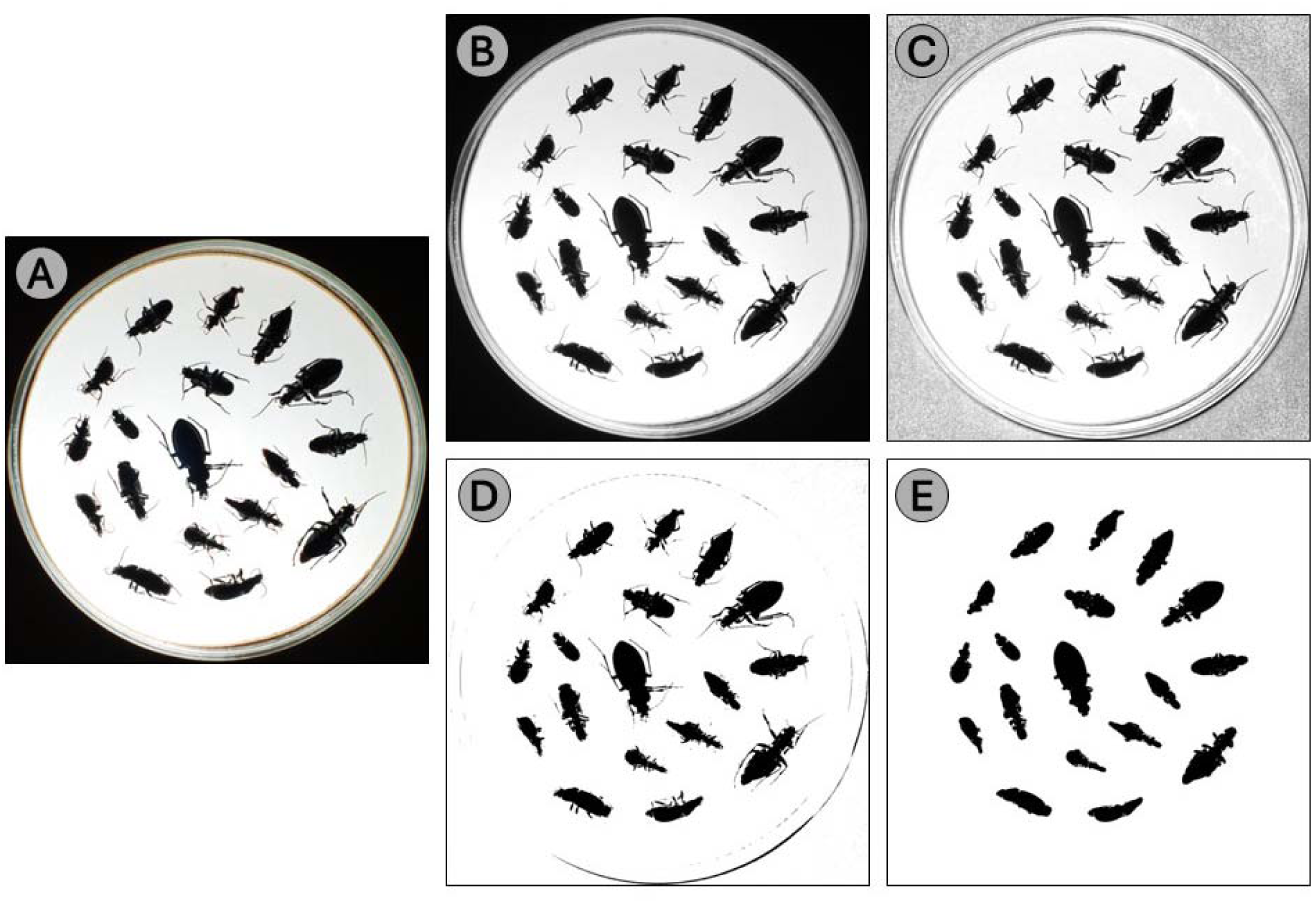
Intermediate images generated during the automated image-processing workflow: A) original; B) grayscale (conversion of the original image to grayscale); C) corrected (image after optional flat-field correction); D) binary (thresholded image separating particles from the background); E) without outliers (binary image after removal of small artefacts and noise).

Additionally, the image-processing step generates four region-of-interest (ROI) files corresponding to different morphometric descriptors (Fig. 3). These include the fitted ellipse (Fig. 3A), the major and minor axes of the fitted ellipse (Fig. 3B), the specimen outline (Fig. 3C), and the maximum and minimum feret diameters (Fig. 3D). Together, these ROI files enable the identification of individual specimens and the quantification of their morphometrics.

**Figure 3.**
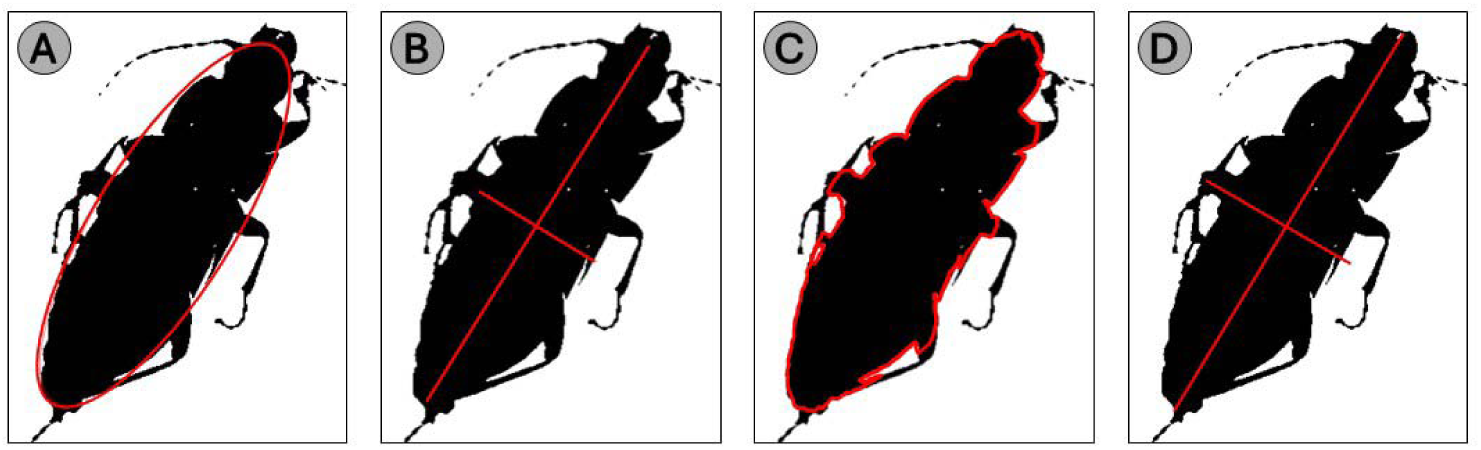
Calculated morphometrics of individual specimen: A) ellipse; B) major and minor ellipse axes; C) outline; D) maximum and minimum ferets.

### 2.4. Application output

At the final stage, AWET generates two CSV output files (Fig. 1J). A regression parameter file, that serves as a repository of the selected allometric models, and a results file that derived from particle analysis outputs in ImageJ. This result file includes additional columns converting pixel-based measurements into calibrated units [mm] for each morphometric variable, as well as estimating fresh body weight [mg] for each detected individual. For a more detailed description of the process, see the user manual in Appendix A4.

## 3. Case example

A standardized image of a Petri dish (Fig. 4) was used as a case example to illustrate the outputs provided by AWET. A total of 18 individuals (carabids in this case) were successfully detected and analyzed. All specimens were processed using identical parameter settings (threshold: 0–120; outlier removal: 25; particle size: 1–50’000; circularity: 0.05–1; scale: 16 pixel mm□¹) and the linear regression model “Carabidae [length]”. For each individual, morphometric parameters including the major and minor axes of the fitted ellipse, maximum and minimum feret diameters, and projected area were extracted (Table 1). Based on these measurements, individual body weight was estimated using two approaches: feret-based and ellipse-based. Estimated individual weights ranged from 73.4 to 700.4 mg for the feret-based approach and from 75.4 to 792.0 mg for the ellipse-based approach. The total biomass of all individuals was 5’848.8 mg and 5’095.0 mg for the feret- and ellipse-based estimates, respectively.

**Figure 4.**
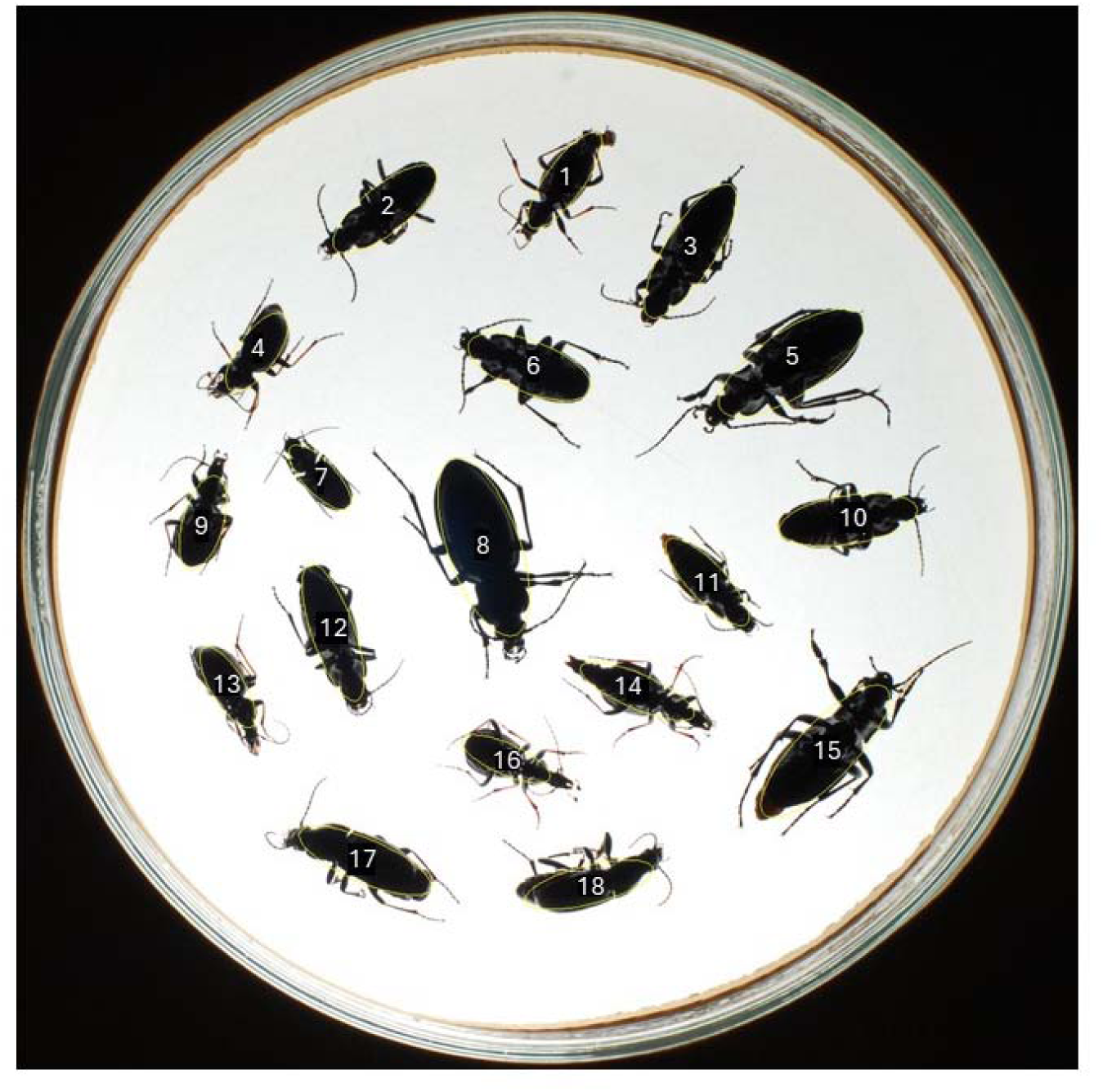
Image of the Petri dish containing 18 carabids used for image-based morphometric analysis. Individuals are labelled (1–18) for correspondence with Table 1.

**Table 1.** Morphometric parameters and derived weight estimates for 18 carabids obtained from image analysis. Recorded measurements include the major and minor axes of the fitted ellipse, maximum and minimum feret diameters, and projected area. Individual body weight was calculated based on feret and ellipse approaches. The sum of each approach shows the total specimen’s biomass of the image.

| ID | Morphometrics |  |  |  |  | Weight estimation |  |
| --- | --- | --- | --- | --- | --- | --- | --- |
|  | Ellipse axe [mm]<br>major | Ellipse axe [mm]<br>minor | Feret [mm]<br>max | Feret [mm]<br>min | Area [mm<sup>2</sup>] | Weight [mg]<br>feret | Weight [mg]<br>ellipse |
| 1 | 16.5 | 5.6 | 18.0 | 6.6 | 72.2 | 240.2 | 188.2 |
| 2 | 17.9 | 6.8 | 18.4 | 8.0 | 94.8 | 255.8 | 233.6 |
| 3 | 21.7 | 7.2 | 22.5 | 8.5 | 122.7 | 443.5 | 404.5 |
| 4 | 13.6 | 5.6 | 14.4 | 6.6 | 59.8 | 126.9 | 109.2 |
| 5 | 24.1 | 9.4 | 25.6 | 11.3 | 177.5 | 643.4 | 543.7 |
| 6 | 18.3 | 7.4 | 18.8 | 8.9 | 106.2 | 269.8 | 248.6 |
| 7 | 11.8 | 4.7 | 11.9 | 5.0 | 43.5 | 75.4 | 73.4 |
| 8 | 26.4 | 10.1 | 27.6 | 11.8 | 209.8 | 792.0 | 700.4 |
| 9 | 13.8 | 6.0 | 14.7 | 7.0 | 65.0 | 135.5 | 113.9 |
| 10 | 19.4 | 7.0 | 20.2 | 8.6 | 107.5 | 328.2 | 295.6 |
| 11 | 16.3 | 5.4 | 17.0 | 6.3 | 69.7 | 204.6 | 182.1 |
| 12 | 20.4 | 6.7 | 20.8 | 7.7 | 107.1 | 356.1 | 338.8 |
| 13 | 13.8 | 5.0 | 14.7 | 5.7 | 54.1 | 134.1 | 112.6 |
| 14 | 17.6 | 5.3 | 20.2 | 7.1 | 73.8 | 328.9 | 224.0 |
| 15 | 24.9 | 8.6 | 26.5 | 10.8 | 168.0 | 704.4 | 589.5 |
| 16 | 13.7 | 5.1 | 14.8 | 6.4 | 54.5 | 137.2 | 111.3 |
| 17 | 20.5 | 6.2 | 20.7 | 6.8 | 100.6 | 352.9 | 346.0 |
| 18 | 19.0 | 5.6 | 20.0 | 6.2 | 83.9 | 319.9 | 279.6 |
|  |  |  |  |  |  | Σ 5'848.8 | Σ 5'095.0 |

## 4. Discussion

AWET is an application that efficiently estimates fresh body weight and extracts morphometric measurements from pre-sorted arthropod samples at the individual level. By combining standardized image analysis with allometric regression models, the software enables rapid, non-destructive quantification of large numbers of specimens using standard laboratory equipment. Compared with traditional biomass estimation methods based on dry or wet weight (e.g. Sage, 1982; Sample et al., 1993), AWET preserves specimens for subsequent taxonomic or molecular analyses. Its reliance on non-expensive imaging equipment and open-source software further provides a favorable cost-benefit ratio, making the approach readily accessible for ecological laboratories and biodiversity monitoring programs.

The primary function of AWET is that it estimates fresh body weight and extracts morphometric measurements for individual specimen, enabling analyses of body-size distributions within and among populations. Individual-level measurements are increasingly recognized as important indicators of ecological processes because body size responds sensitively to changes in environmental conditions and resource availability (Gardner et al., 2011; Tseng et al., 2018). Consequently, AWET not only facilitates biomass estimation but also provides a valuable source of functional trait data that can be integrated into ecological and evolutionary analyses.

Image recognition and machine learning have become valuable tools for entomologists (Høye et al., 2021; Sys et al., 2022; van Klink et al., 2022). Several automated image-analysis frameworks for arthropods have recently been developed, but these occupy different operational niches (Table 2). Some frameworks, such as InsectMorphoAI (Shirali et al., 2026), focus on detailed morphometric analysis and biomass estimation using deep-learning-based segmentation models. Others, including DiversityScanner (Wührl et al., 2022) and BIODISCOVER (Ärje et al., 2020), combine automated imaging with robotic specimen handling and machine-learning-based identification to maximize throughput, albeit at the cost of dedicated hardware and specialized imaging platforms. The tool from Schneider et al. (2022) similarly estimates biomass from images and relies on trained classification models and predefined taxonomic categories.

**Table 2.** Overview of the main characteristics of AWET and other existing image-based arthropod analysis tools.

| Characteristics | AWET<br>(this study) | InsectMorphoAI<br>(Shirali et al., 2026) | DiversityScanner<br>(Wühl et al., 2022) | BIODISCOVER<br>(Ärje et al., 2020) | PracticalTool<br>(Schneider et al., 2022) |
| --- | --- | --- | --- | --- | --- |
| Primary function | Arthropod fresh body weight estimation | Arthropod biomass estimation | Insect sorting | Arthropod imaging and identification | Arthropod classification |
| Secondary output | Morphometric measurements | Morphometric measurements<br>Volume estimation | Morphometric measurements | Biomass estimation | Biomass estimation |
| Biomass type | Fresh weight | Dry and wet weight | Not available | Dry weight | Class-specific biomass |
| Morphometrics | Length, width, area, weight | Length, width, volume | Length, volume | Area, weight | Area, weight |
| Abundance estimation | Yes | Yes | Yes | Yes | Yes |
| Measurement level | Specimen | Specimen | Specimen | Specimen | Specimen |
| Dimensionality | 2D | 2D (3D) | 2D | 2D | 2D |
| Pre-sorting required | Yes | No | No | No | Yes |
| Taxonomic range | Up to 124 taxonomic groups | Up to 3 taxonomic groups | Up to 14 taxa (<3 mm) | Up to 3 taxonomic groups and 12 taxa<br>(small enough for the funnel) | Up to 18 taxonomic groups |
| Non-destructive | Yes | Yes | Yes | Yes | Yes |
| Processing speed | 44 ms/specimen | 54'000 ms/specimen | 13'000 ms/specimen | 6'000 ms/specimen | Not specified |
| Machine/deep learning required | No | Yes | Yes | Yes | Yes |
| Flexibility | Can be adapted to specific taxonomic groups | Requires a trained dataset | Requires a trained dataset | Requires a trained dataset | Requires a trained dataset |
| Software | Open source | Open source | Open source | Open source | Open source |
| Hardware | Standard equipment<br>(< €1,000) | ENTOMOSCOPE imaging system<br>(< €1,000) | Specific imaging platform<br>(< €5,000) | BIODISCOVER imaging system<br>(~ €5,000) | Standard equipment<br>(< €1,000) |

AWET occupies a complementary niche by emphasizing accessibility, computational efficiency, and rapid fresh body weight estimation without requiring machine-learning models or specialized imaging systems. With an average processing time of approximately 44 ms per specimen, AWET substantially exceeds the processing speeds reported for comparable image-based approaches (Table 2) while remaining deployable and affordable using standard imaging equipment. One of the principal strengths of AWET is its taxonomic flexibility.

Unlike machine-learning-based frameworks, whose applicability is constrained by the taxonomic groups represented in their training datasets, AWET does not depend on trained classification or segmentation models. Instead, users can select the taxonomic grouping that best matches their study objectives and available reference data. Body-weight estimation can therefore be performed for each specimen at the available order or family levels, allowing researchers to trade off estimation accuracy against sample size and taxonomic certainty. The current application includes allometric regression models for 25 taxonomic groups (order and family level) and can be expanded to as many as 124 groups (family level). Importantly, this does not represent a fixed limit of the framework. Because AWET is based on transparent statistical models rather than deep-learning classifiers, additional taxonomic groups can be incorporated simply by generating suitable reference measurements and fitting new allometric regression models without modifying the underlying software architecture. This flexibility makes the framework readily transferable to different geographic regions, arthropod communities, and long-term monitoring programs where taxonomic resolution often varies among projects.

Despite these advantages, several limitations should be considered. AWET requires specimens to be sorted into taxonomic groups before proceeding because body weight estimation relies on taxon-specific allometric regression models rather than automated taxonomic recognition. However, this sorting step is usually already part of ecological monitoring workflows in order to assess biodiversity. Furthermore, body weight estimates for taxa such as Lepidoptera should be interpreted with caution because variable wing positions may influence image-derived morphometric measurements and consequently increase estimation uncertainty. Similarly, Feret-based measurements may slightly overestimate body dimensions compared with ellipse-based measurements, which more closely approximate true body shape. Finally, estimation accuracy depends on the representativeness of the calibration data used to develop statistical models. Although logarithmic transformation reduces the influence of extreme values, estimation uncertainty may increase when specimens fall outside the size range represented in the reference dataset. Future refinement of the underlying models through additional calibration data will further improve estimation accuracy and extend the taxonomic coverage of the framework.

## 5. Conclusion

Overall, AWET is a practical and scalable solution for image-based arthropod biomass estimation that complements existing automated phenotype approaches. By combining rapid processing, individual-level morphometric information, flexible taxonomic grouping, and non-destructive analysis within an accessible workflow, the open-source software bridges the gap between traditional, laborious biomass measurements and highly specialized machine-learning-based systems. As standardized imaging becomes increasingly integrated into biodiversity monitoring, AWET has the potential to become an important component of ecological workflows, enabling reproducible quantification of arthropod biomass, body-size, and functional traits across broad spatial and temporal scales.

## Acknowledgements

We would like to acknowledge the contributions of Lucas Marian, Esra Sohlström, Andrew D. Barnes, Noor F. Haneda, Stefan Scheu, Björn C. Rall and Ulrich Brose in obtaining the Sohlström et al. (2018) measurements that provided the data basis and enabled the image recognition tools that has led to AWET.

## Author contributions

MC, JB, MKO and NvK conceptualized the paper, defining the overarching research goals and aims. Methodology was developed by MC, JB and MKO. Software development was performed by TZ, RB, MC and MKO. Data curation and formal analysis for regression models were carried out by RF and MC. Investigation, including acquisition of standardized images of pre-sorted arthropod specimens, was performed by MC, MKO and NvK. Validation of the workflow and body weight estimation outputs was undertaken by MC and MKO. Visualization of figures and workflow diagrams was performed by MC. Resources, including provision of datasets, imaging infrastructure, and computational tools, were provided by MKO. Supervision and project administration were led by MC, ensuring coordination and oversight of the research activities. JB and MKO prepared the original draft of the manuscript, while all authors contributed to writing, reviewing, and editing. Funding acquisition was secured by KB and MKO. All authors approved the final version of the manuscript.

## Application availability

Cretton, M., Boesch, R., Bolliger, J., Bollmann, K., Flury, R., Jochum, M., van Koppenhagen, N., Zimmermann, T., Obrist, M. K. (2026). AWET - Arthropod Weight Estimation Tool. *EnviDat.* https://www.doi.org/10.16904/envidat.771.

## Blind review of application availability (EnviDat)

https://www.envidat.ch/#/review/c8ee7a59-d7ff-45b0-b819-ed27f9b57c43.

## Application history

https://gitlabext.wsl.ch/cretton/awet.

---

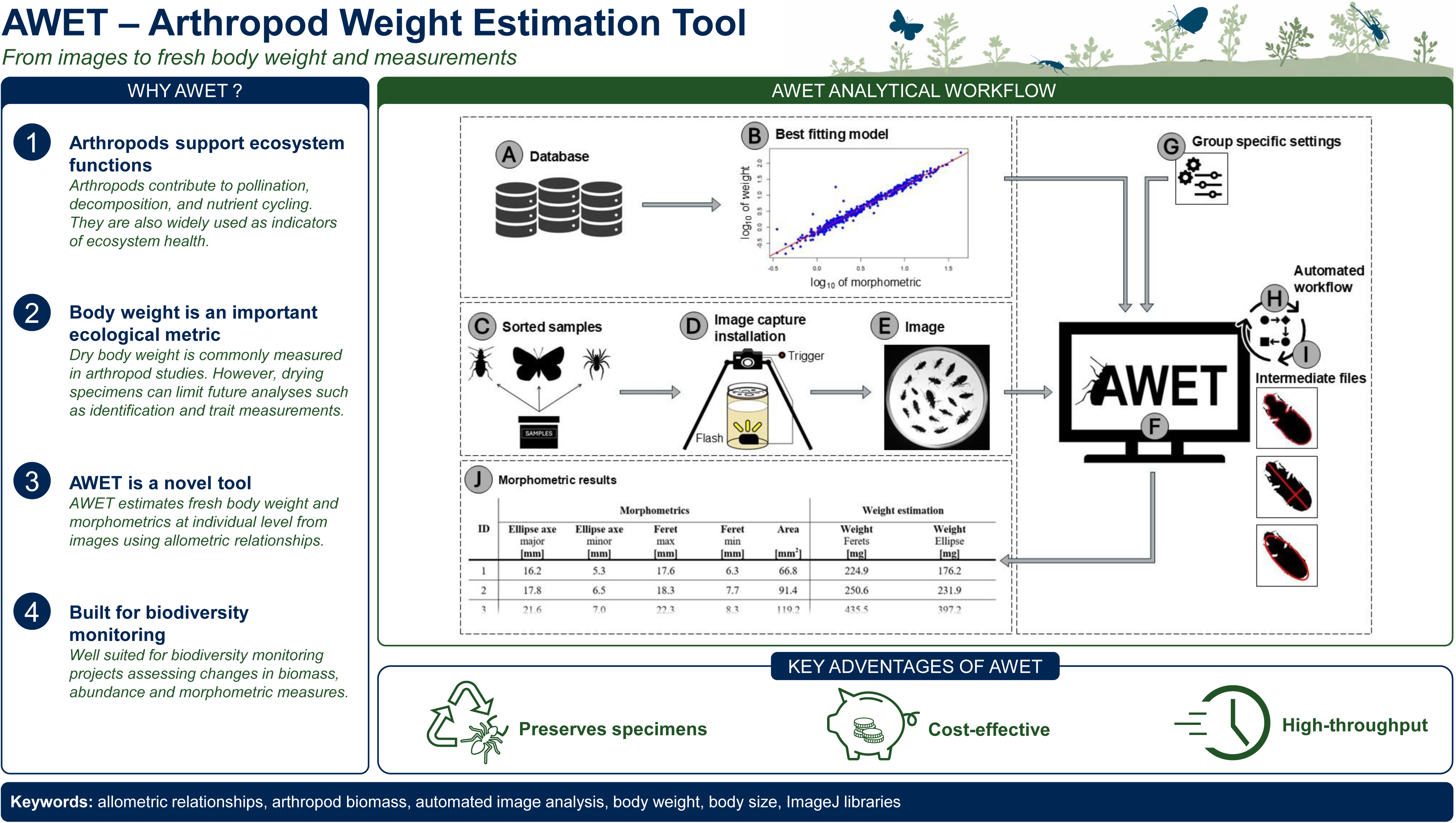

