## supplement file for "AWET – Arthropod Weight Estimation Tool"

### A1 – Classification of arthropods in different taxonomic groups

Table A1.1: Classification in class(es), order(s), group(s) and the included families.

| **Class(es)** | **Orders(s) – Group(s)** | **Included families** |
| --- | --- | --- |
| Arachnida | Araneae | Agelenidae, Amaurobiidae, Anyphaenidae, Araneidae, Clubionidae, Dysderidae, Gnaphosidae, Linyphiidae, Liocranidae, Lycosidae, Philodromidae, Pholcidae, Pisauridae, Salticidae, Tetragnathidae, Thomisidae |
| Arachnida | Opiliones and Pseudoscorpionida | Larcidae, Nemastomatidae, Neobisiidae, Phalangiidae, Trogulidae |
| Insecta | Coleoptera I  *(Carabidae)* | Carabidae |
| Insecta | Coleoptera II  *(Staphylinidae)* | Staphylinidae |
| Insecta | Coleoptera III *(sapro-xylophagous)* | Anobiidae, Cantharidae, Curculionidae, Elateridae, Endomychidae, Melyridae, Nitidulidae, Oedemeridae |
| Insecta | Coleoptera Rest I *(without Carabidae and Staphylinidae)* | Anobiidae, Cantharidae, Chrysomelidae, Coccinellidae, Curculionidae, Dascillidae, Elateridae, Endomychidae, Leiodidae, Melyridae, Nitidulidae, Oedemeridae, Tenebrionidae |
| Insecta | Coleoptera Rest II *(without Carabidae, Staphylinidae and sapro-xylophagous)* | Chrysomelidae, Coccinellidae, Dascillidae, Leiodidae |
| Insecta | Coleoptera All | Anobiidae, Cantharidae, Carabidae, Chrysomelidae, Coccinellidae, Curculionidae, Dascillidae, Elateridae, Endomychidae, Leiodidae, Melyridae, Nitidulidae, Oedemeridae, Staphylinidae, Tenebrionidae |
| Insecta | Dermaptera | Forficulidae, Labiidae |
| Insecta | Diptera I  *(Syrphidae)* | Syrphidae |
| Insecta | Diptera All | Anthomyiidae, Asilidae, Athericidae, Canacidae, Cecidomyiidae, Chaoboridae, Chloropidae, Conopidae, Culicidae, Dolichopodidae, Dryomyzidae, Empididae, Fanniidae, Lauxaniidae, Lonchopteridae, Milichiidae, Muscidae, Periscelididae, Phoridae, Pipunculidae, Platypezidae, Psilidae, Rhagionidae, Scathophagidae, Sciaridae, Sciomyzidae, Sepsidae, Stratiomyidae, Syrphidae, Therevidae, Tipulidae, Dip_Unknown |
| Insecta | Heteroptera | Lygaeidae, Microphysidae, Miridae, Nabidae, Pentatomidae, Pyrrhocoridae, Rhopalidae, Rhyparochromidae, Saldidae |
| Insecta | Homoptera | Aphididae, Aphrophoridae, Cicadellidae, Delphacidae, Issidae |
| Insecta | Hymenoptera I *(Apocrita/Aculeata)* | Apidae, Bethylidae, Formicidae, Vespidae |
| Insecta | Hymenoptera II  *(Formicidae)* | Formicidae |
| Insecta | Hymenoptera III *(Apocrita/Terebrantes)* | Braconidae, Chalcididae, Diapriidae, Eulophidae, Ichneumonidae, Pamphiliidae, Proctotrupidae, Pteromalidae, Scelionidae, Tenthredinidae |
| Insecta | Hymenoptera All | Apidae, Bethylidae, Braconidae, Chalcididae, Diapriidae, Eulophidae, Formicidae, Ichneumonidae, Pamphiliidae, Proctotrupidae, Pteromalidae, Scelionidae, Tenthredinidae, Vespidae |
| Insecta | Lepidoptera | Hesperiidae, Lycaenidae, Ypsolophidae, Zygaenidae, Lep_Unknown |
| Insecta | Neuroptera and Mecoptera | Chrysopidae, Hemerobiidae, Panorpidae |
| Insecta | Orthoptera | Acrididae, Tettigoniidae |
| Malacostraca | Isopoda | Cylisticidae, Ligiidae, Oniscidae, Philosciidae, Porcellionidae, Trichoniscidae, Iso_Unknown |
| Chilopoda | Lithobiomorpha | Lithobiidae |
| Diplopoda | Glomerida | Glomeridae |
| Diplopoda | Chordeumatida and Julida | Chordeumatoidae, Julidae |
| Diplopoda | Polydesmida | Polydesmidae |

### A2 – Allometric regression models of each taxonomic group

Table A2.1: Allometric regression models resulting from the temperate arthropod dataset (Sohlström et al. (2018). The models are classified in classes and taxonomic order groups (Appendix A1, Table A1.1). The models contain the intercept and the allometric coefficients: log_10_(width_max_) is the slope of the maximal width; log_10_(length) is the slope of the length; log_10_(area) the slope of the area – (***p < 0.001; **p < 0.01; *p < 0.05). N is the number of individuals included in the regression models. For each model Akaike Information Criterion (AIC), Bayesian Information Criterion (BIC), coefficient of determination ($R^{2}$), adjusted $R^{2}$ and the root mean squared error (RMSE) are reported. In each taxonomic group, the models are sorted according to the BIC.

| **Classes** | **Taxonomic groups** | **Intercept** | | **log_10_(width_max_)** | | **log_10_(length)** | | **log_10_(area)** | | **N** | **AIC** | **BIC** | **R^2^** | **adj. R^2^** | **RMSE** |
| --- | --- | --- | --- | --- | --- | --- | --- | --- | --- | --- | --- | --- | --- | --- | --- |
| Arachnida | Araneae | -0.123 | *** |  |  |  |  | 1.427 | *** | 519 | -930.62 | -917.87 | 0.97 | 0.97 | 0.10 |
|  |  | -0.733 | *** |  |  | 2.623 | *** |  |  | 519 | -485.09 | -472.34 | 0.93 | 0.93 | 0.15 |
|  |  | 0.243 | *** | 2.853 | *** |  |  |  |  | 519 | -481.46 | -468.71 | 0.93 | 0.93 | 0.15 |
|  | Opiliones and Pseudoscorpionida | -0.382 | *** |  |  | 2.390 | *** |  |  | 94 | -48.38 | -40.75 | 0.90 | 0.90 | 0.18 |
|  |  | 0.027 |  | 2.872 | *** |  |  |  |  | 94 | -29.88 | -22.25 | 0.88 | 0.88 | 0.20 |
|  |  | -0.523 | *** |  |  |  |  | 1.259 | *** | 94 | -14.33 | -6.70 | 0.86 | 0.85 | 0.22 |
| Insecta | Carabidae | -1.140 | *** |  |  | 2.802 | *** |  |  | 111 | -202.20 | -194.07 | 0.91 | 0.91 | 0.09 |
|  |  | -1.076 | *** | 0.144 |  | 2.657 | *** |  |  | 111 | -203.03 | -192.19 | 0.91 | 0.91 | 0.09 |
|  |  | -0.910 | *** |  |  |  |  | 1.675 | *** | 111 | -97.32 | -89.19 | 0.76 | 0.76 | 0.15 |
|  |  | 1.057 | *** | 1.498 | *** |  |  |  |  | 111 | -17.16 | -9.03 | 0.51 | 0.51 | 0.22 |
|  | Staphylinidae | 0.175 | *** | 2.942 | *** |  |  |  |  | 55 | -49.45 | -43.43 | 0.97 | 0.97 | 0.15 |
|  |  | -1.578 | *** |  |  | 2.635 | *** |  |  | 55 | -29.58 | -23.56 | 0.95 | 0.95 | 0.18 |
|  |  | -1.920 | *** |  |  |  |  | 2.107 | *** | 55 | 34.15 | 40.17 | 0.85 | 0.85 | 0.31 |
|  | Coleoptera *sapro-xylobiont* | 0.151 | *** | 2.894 | *** |  |  |  |  | 104 | -117.53 | -109.59 | 0.94 | 0.94 | 0.13 |
|  |  | -0.835 | *** |  |  | 2.275 | *** |  |  | 104 | -113.08 | -105.15 | 0.94 | 0.94 | 0.14 |
|  |  | -0.690 | *** |  |  |  |  | 1.109 | *** | 104 | -76.83 | -68.90 | 0.91 | 0.91 | 0.16 |
|  | Coleoptera *without carab. & staph.* | -0.406 | *** | 1.378 | *** | 1.253 | *** |  |  | 216 | -477.73 | -464.23 | 0.97 | 0.97 | 0.08 |
|  |  | -0.706 | *** |  |  | 2.202 | *** |  |  | 216 | -197.96 | -187.83 | 0.88 | 0.88 | 0.15 |
|  |  | 0.116 | *** | 2.560 | *** |  |  |  |  | 216 | -178.22 | -168.09 | 0.87 | 0.87 | 0.16 |
|  |  | -0.319 | *** |  |  |  |  | 0.875 | *** | 216 | 52.67 | 62.79 | 0.62 | 0.61 | 0.27 |
|  | Coleoptera *without carab. & staph. & sapro.* | -0.420 | *** | 1.218 | *** | 1.344 | *** |  |  | 110 | -261.83 | -251.03 | 0.94 | 0.94 | 0.07 |
|  |  | -0.692 | *** |  |  | 2.319 | *** |  |  | 110 | -167.08 | -158.98 | 0.85 | 0.85 | 0.11 |
|  |  | 0.105 | *** | 2.269 | *** |  |  |  |  | 110 | -155.52 | -147.42 | 0.83 | 0.83 | 0.12 |
|  |  | 0.252 | * |  |  |  |  | 0.392 | *** | 110 | 23.27 | 31.37 | 0.13 | 0.13 | 0.26 |
|  | Coleoptera *all* | 0.139 | *** | 2.662 | *** |  |  |  |  | 382 | -106.08 | -94.24 | 0.94 | 0.94 | 0.21 |
|  |  | -0.938 | *** |  |  | 2.501 | *** |  |  | 382 | 69.53 | 81.37 | 0.90 | 0.90 | 0.26 |
|  |  | -1.013 | *** |  |  |  |  | 1.589 | *** | 382 | 342.48 | 354.31 | 0.80 | 0.80 | 0.38 |
|  | Dermaptera | -1.093 | *** |  |  |  |  | 1.340 | *** | 60 | -105.73 | -99.45 | 0.97 | 0.97 | 0.10 |
|  |  | 0.235 | *** | 3.117 | *** |  |  |  |  | 60 | -86.11 | -79.83 | 0.95 | 0.95 | 0.11 |
|  |  | -0.947 | *** |  |  | 2.337 | *** |  |  | 60 | -86.00 | -79.72 | 0.95 | 0.95 | 0.11 |
|  | Syrphidae | -0.026 |  | 1.994 | *** | 0.480 | * |  |  | 124 | -192.31 | -181.02 | 0.68 | 0.68 | 0.11 |
|  |  | 0.278 | *** | 2.343 | *** |  |  |  |  | 124 | -187.76 | -179.30 | 0.67 | 0.66 | 0.11 |
|  |  | -0.484 | ** |  |  | 1.756 | *** |  |  | 124 | -119.91 | -111.45 | 0.43 | 0.42 | 0.15 |
|  |  | 0.847 | *** |  |  |  |  | 0.151 | * | 124 | -57.93 | -49.46 | 0.05 | 0.05 | 0.19 |
|  | Diptera *all* | -0.309 | *** | 1.595 | *** | 0.997 | *** |  |  | 504 | -707.58 | -690.69 | 0.94 | 0.94 | 0.12 |
|  |  | 0.236 | *** | 2.425 | *** |  |  |  |  | 504 | -515.66 | -502.99 | 0.91 | 0.91 | 0.14 |
|  |  | -1.057 | *** |  |  | 2.489 | *** |  |  | 504 | -279.39 | -266.72 | 0.85 | 0.85 | 0.18 |
|  |  | -0.215 | *** |  |  |  |  | 0.837 | *** | 504 | 380.91 | 393.58 | 0.44 | 0.44 | 0.35 |
|  | Heteroptera | -0.430 | *** | 1.425 | *** | 1.186 | *** |  |  | 437 | -1009.11 | -992.79 | 0.96 | 0.96 | 0.08 |
|  |  | -0.665 | *** |  |  |  |  | 1.182 | *** | 437 | -439.24 | -427.00 | 0.86 | 0.86 | 0.15 |
|  |  | 0.342 | *** | 1.886 | *** |  |  |  |  | 437 | -419.09 | -406.85 | 0.86 | 0.85 | 0.15 |
|  |  | -0.901 | *** |  |  | 2.368 | *** |  |  | 437 | -11.74 | 0.50 | 0.63 | 0.63 | 0.24 |
|  | Homoptera | -0.895 | *** |  |  |  |  | 1.346 | *** | 140 | -316.29 | -307.47 | 0.96 | 0.96 | 0.08 |
|  |  | 0.116 | *** | 2.591 | *** |  |  |  |  | 140 | -292.26 | -283.44 | 0.96 | 0.96 | 0.08 |
|  |  | -1.041 | *** |  |  | 2.619 | *** |  |  | 140 | -203.49 | -194.67 | 0.92 | 0.92 | 0.11 |
|  | Aculeata | 0.174 | *** | 2.627 | *** |  |  |  |  | 141 | -211.72 | -202.87 | 0.96 | 0.96 | 0.11 |
|  |  | -1.553 | *** |  |  | 3.157 | *** |  |  | 141 | -115.77 | -106.93 | 0.92 | 0.92 | 0.16 |
|  |  | -1.139 | *** |  |  |  |  | 1.632 | *** | 141 | 103.83 | 112.68 | 0.63 | 0.63 | 0.34 |
|  | Formicidae | -0.618 | *** | 1.794 | *** | 1.367 | *** |  |  | 98 | -183.96 | -173.62 | 0.85 | 0.85 | 0.09 |
|  |  | 0.179 | *** | 2.537 | *** |  |  |  |  | 98 | -138.31 | -130.55 | 0.76 | 0.76 | 0.12 |
|  |  | -1.309 | *** |  |  | 2.744 | *** |  |  | 98 | -95.29 | -87.53 | 0.63 | 0.63 | 0.14 |
|  |  | -0.568 | *** |  |  |  |  | 0.932 | *** | 98 | -82.87 | -75.11 | 0.58 | 0.58 | 0.15 |
|  | Hymenoptera *without acule.* | -0.351 | ** | 1.931 | *** | 1.015 | *** |  |  | 81 | -74.19 | -64.62 | 0.92 | 0.92 | 0.15 |
|  |  | -0.453 | *** |  |  |  |  | 1.459 | *** | 81 | -60.26 | -53.08 | 0.91 | 0.91 | 0.16 |
|  |  | 0.254 | *** | 2.876 | *** |  |  |  |  | 81 | -48.71 | -41.52 | 0.89 | 0.89 | 0.17 |
|  |  | -1.285 | *** |  |  | 2.583 | *** |  |  | 81 | -9.64 | -2.46 | 0.82 | 0.82 | 0.22 |
|  | Hymenoptera *all* | 0.207 | *** | 2.600 | *** |  |  |  |  | 222 | -222.31 | -212.10 | 0.94 | 0.94 | 0.14 |
|  |  | -1.486 | *** |  |  | 3.018 | *** |  |  | 222 | -90.12 | -79.91 | 0.89 | 0.89 | 0.19 |
|  |  | -0.447 | *** |  |  |  |  | 1.112 | *** | 222 | 159.22 | 169.43 | 0.67 | 0.67 | 0.34 |
|  | Lepidoptera | -0.499 | *** |  |  |  |  | 1.401 | *** | 29 | -48.16 | -44.06 | 0.96 | 0.95 | 0.10 |
|  |  | 0.227 | *** | 2.882 | *** |  |  |  |  | 29 | -44.14 | -40.04 | 0.95 | 0.95 | 0.10 |
|  |  | -1.274 | *** |  |  | 2.505 | *** |  |  | 29 | -20.72 | -16.61 | 0.88 | 0.88 | 0.15 |
|  | Neuroptera and Mecoptera | 0.435 | *** | 3.121 | *** |  |  |  |  | 28 | -38.12 | -34.12 | 0.89 | 0.89 | 0.11 |
|  |  | 0.568 | * | 3.230 | *** | -0.158 |  |  |  | 28 | -36.41 | -31.08 | 0.90 | 0.89 | 0.11 |
|  |  | -0.614 | ** |  |  |  |  | 1.583 | *** | 28 | -19.69 | -15.70 | 0.80 | 0.79 | 0.15 |
|  |  | -1.023 |  |  |  | 2.190 | *** |  |  | 28 | 9.79 | 13.79 | 0.42 | 0.40 | 0.26 |
|  | Orthoptera | -0.142 | * |  |  |  |  | 1.334 | *** | 35 | -84.35 | -79.69 | 0.97 | 0.97 | 0.07 |
|  |  | 0.136 |  | 1.713 | *** | 0.823 | *** |  |  | 35 | -70.05 | -63.83 | 0.95 | 0.95 | 0.08 |
|  |  | 0.724 | *** | 2.417 | *** |  |  |  |  | 35 | -54.15 | -49.48 | 0.92 | 0.92 | 0.10 |
|  |  | -0.640 | ** |  |  | 2.267 | *** |  |  | 35 | -24.83 | -20.17 | 0.83 | 0.82 | 0.16 |
| Chilopoda | Lithobiomorpha | -1.671 | *** |  |  | 2.780 | *** |  |  | 161 | -275.19 | -265.95 | 0.97 | 0.97 | 0.10 |
|  |  | 0.628 | *** | 3.043 | *** |  |  |  |  | 161 | -274.39 | -265.15 | 0.97 | 0.97 | 0.10 |
|  |  | -1.139 | *** |  |  |  |  | 1.580 | *** | 161 | -56.61 | -47.36 | 0.90 | 0.90 | 0.20 |
| Diplopoda | Chordeumatida and Julida | -1.784 | *** |  |  | 2.591 | *** |  |  | 131 | -211.56 | -202.94 | 0.93 | 0.93 | 0.11 |
|  | Glomerida | -0.747 | *** |  |  | 2.510 | *** |  |  | 182 | -260.01 | -250.40 | 0.87 | 0.87 | 0.12 |
|  | Polydesmida | -1.400 | *** |  |  | 2.500 | *** |  |  | 37 | -79.35 | -74.52 | 0.86 | 0.85 | 0.08 |
|  |  | -1.400 | *** | 0.215 |  | 2.443 | *** |  |  | 12 | -39.11 | -37.17 | 0.98 | 0.98 | 0.03 |
|  |  | 0.834 | *** | 2.452 | ** |  |  |  |  | 12 | -5.51 | -4.06 | 0.68 | 0.64 | 0.15 |
|  |  | 0.473 |  |  |  |  |  | 0.693 |  | 12 | 3.34 | 4.79 | 0.32 | 0.26 | 0.22 |
| Malacostraca | Isopoda | -0.070 | ** | 2.498 | *** |  |  |  |  | 88 | -185.07 | -177.64 | 0.98 | 0.98 | 0.08 |
|  |  | -1.292 | *** |  |  | 2.950 | *** |  |  | 88 | -133.61 | -126.18 | 0.97 | 0.97 | 0.11 |
|  |  | -1.121 | *** |  |  |  |  | 1.284 | *** | 88 | -14.92 | -7.48 | 0.88 | 0.88 | 0.21 |

### A3 – Validation of image-based weight estimation methods

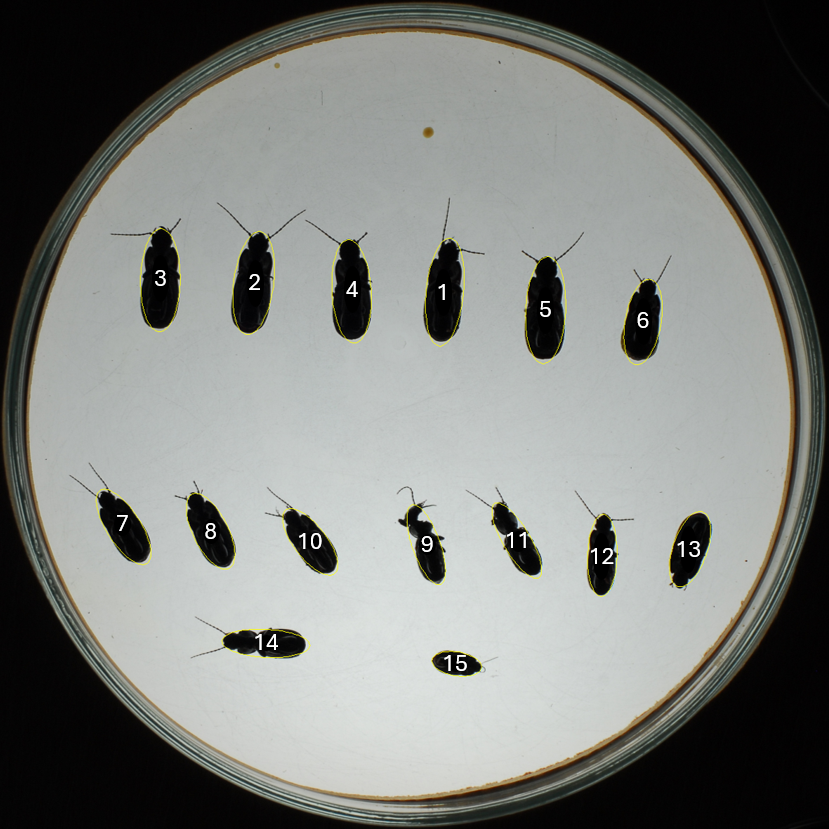

Figure A3.1: Image of the Petri dish containing 15 carabids used for image-based morphometric analysis. Individuals are labelled (1–15) for correspondence with Table A3.1.

Table A3.1: Effective fresh body weight of 15 carabids measured with a precision balance and fresh weight estimations obtained from image analysis from Fig. A3.1, using the four linear regression models. Individual body weight was calculated based on feret and ellipse approaches.

|  | **Weight effective** *[mg]* |  | **Weight estimate** *[mg]* | | | |
| --- | --- | --- | --- | --- | --- | --- |
| **Beetle ID** |  | **Model** | **length** | **length & width** | **area** | **width** |
| **1** | 330 |  | 291 | 287 | 277 | 194 |
| **2** | 290 |  | 259 | 260 | 292 | 216 |
| **3** | 300 |  | 253 | 255 | 285 | 213 |
| **4** | 290 |  | 255 | 256 | 281 | 210 |
| **5** | 280 |  | 280 | 281 | 306 | 215 |
| **6** | 230 |  | 155 | 158 | 180 | 184 |
| **7** | 180 |  | 155 | 156 | 162 | 167 |
| **8** | 180 |  | 137 | 140 | 160 | 176 |
| **9** | 170 |  | 144 | 142 | 112 | 125 |
| **10** | 140 |  | 125 | 126 | 134 | 159 |
| **11** | 140 |  | 153 | 150 | 112 | 121 |
| **12** | 170 |  | 133 | 133 | 118 | 137 |
| **13** | 140 |  | 118 | 121 | 143 | 173 |
| **14** | 160 |  | 162 | 159 | 122 | 127 |
| **15** | 50 |  | 33 | 34 | 36 | 100 |

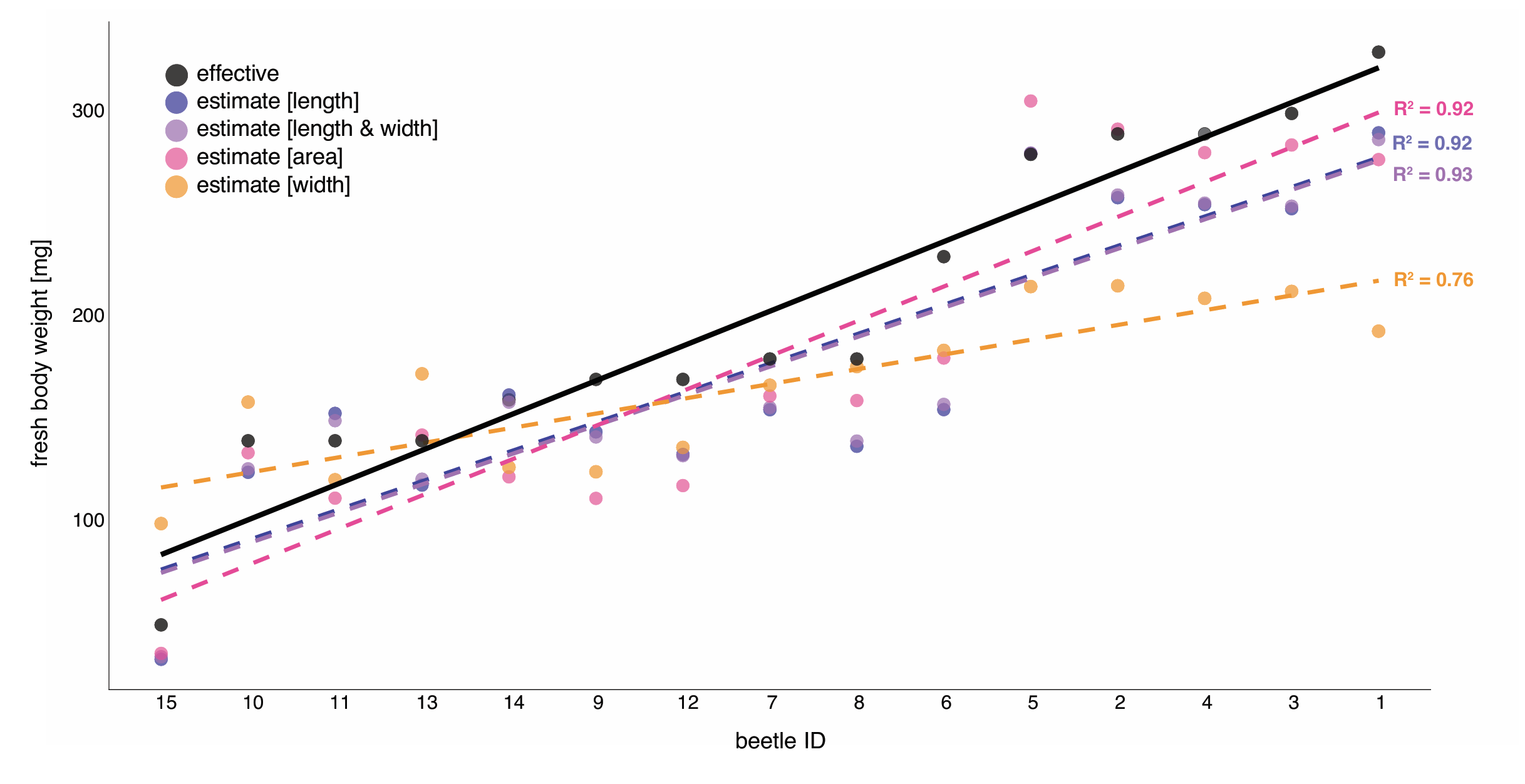

Figure A3.2: Overview of effective fresh body weight (black) of the 15 carabids from Fig. A3.1 and the respective results of the four linear regression models for fresh body weight. Estimation model with “length” (blue), “area” (pink), “width” (orange), and with the combination of “length and width” (purple).

### A4 – Manual use of AWET

The tool AWET (Arthropod Weight Estimation Tool) can measure various morphometric variables of individual arthropod samples using image analysis. AWET is a Java application built around ImageJ libraries and is designed to measure length, width and area and to calculate body mass of individual arthropods from images of specimens. For individual morphometrics and body mass calculation, specimens need to be sorted into pre-defined taxonomic groups.

With his graphical user interface, AWET offers a variety of user selectable options and settings (Fig. A4.1):

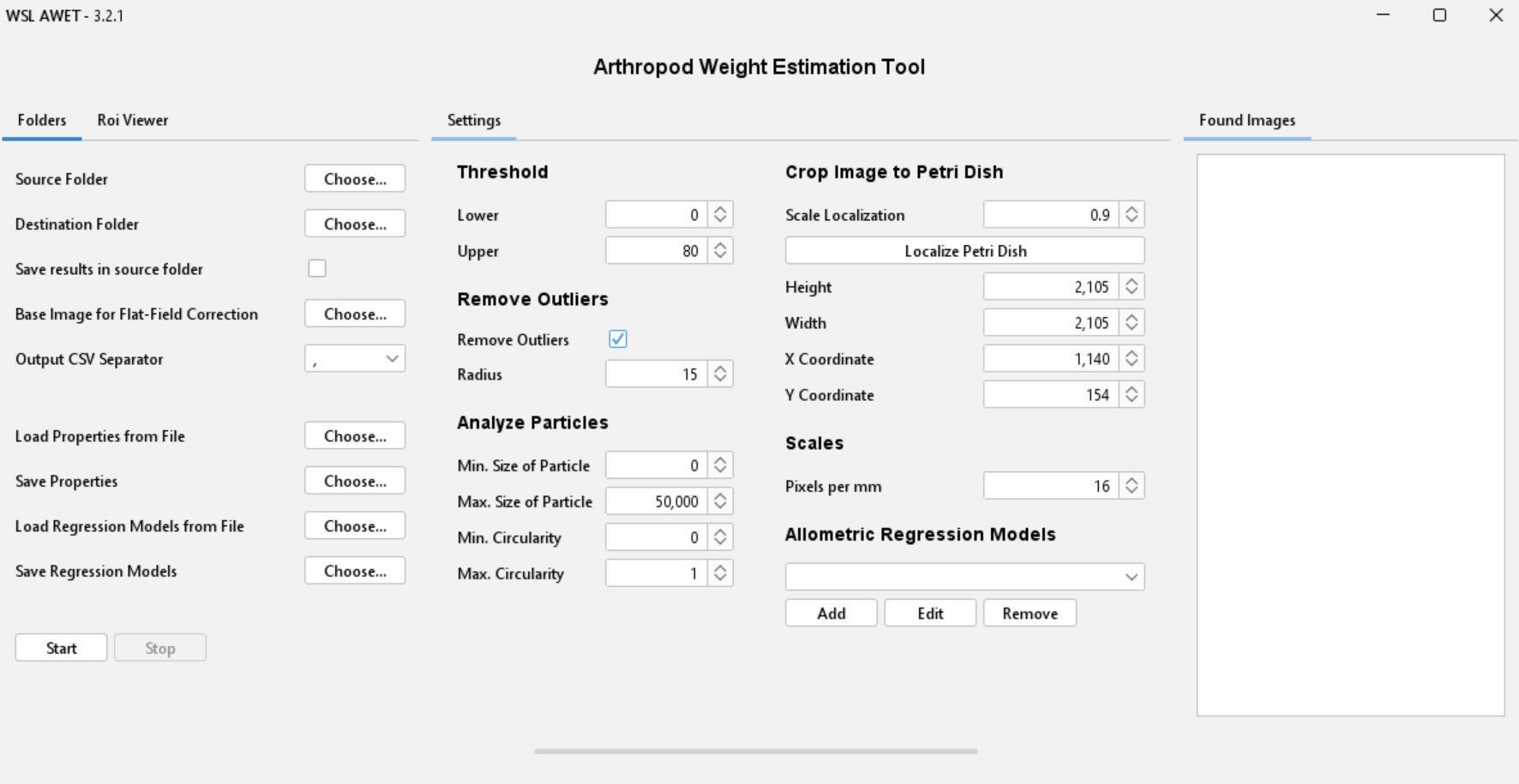

Figure A4.1: Graphical user interface for AWET.

**AWET Preparation – Input and Output (in Folders tab)**

**Source Folder**

Select the directory containing the images to be analyzed, the drag and drop is available. The selected directory and all its subdirectories will be scanned for (.jpg) images. All detected images are listed under the Found Images tab.

**Destination Folder**

Select the directory where all output files and results will be stored.

**Save Results in Source Folder**

When this option is enabled, the destination folder is automatically set to the same directory as the source folder.

**Base Image for Flat-Field Correction**

Select an image captured using the same camera setup as the source images but without specimens present. This image is used for flat-field correction to remove illumination gradients and camera artefacts from the analyzed images.
If no such image is provided, the correction step is skipped. However, this may make it more difficult to determine suitable threshold settings for reliable object detection.

**Output CSV Separator**

Define the separator used in the generated result files. Available options are:

- TAB (tabulation)
- ; (semicolon)
- , (comma)

**AWET Preparation – Models and Properties (in Folders tab)**

**Load Properties from File**

Load previously saved software settings from a properties file into the settings panel.

**Save Properties**

Save the current settings from the settings panel to a properties file for later reuse. Note that allometric regression models are not stored in this file and must be saved separately.

**Load Regression Models from File**

Load previously defined regression models from a .csv file with the following structure:

Table A4.1: Structure of the regression model csv-file.

| **Column** | **Description** |
| --- | --- |
| Title | Unique identifier of the regression model |
| Region | Geographic region |
| Taxonomic Group | Arthropod group used for the model |
| Intercept | Regression intercept |
| Slope Length | Coefficient for specimen length |
| Slope Width | Coefficient for specimen width |
| Slope Area | Coefficient for specimen area |

The Title must be unique, as it is used by the application to identify each model.

**Save Regression Models**

Export all currently loaded regression models to a .csv file.

**AWET Settings (in Settings tab)**

**Threshold**

Threshold values are used to generate the binary image required for object detection. Pixel values within the defined lower and upper threshold range are set to black, while all others are set to white.

Both threshold values can range between 0 and 255.

**Remove Outliers**

This optional step removes small artefacts such as dirt particles or detached arthropod fragments (e.g. legs or wings) that may remain after thresholding.

The procedure replaces each pixel with the median value of surrounding pixels within a defined radius. Because the operation is applied to a binary image, the resulting pixels remain either black or white.

This step can also help to reconnect segmented arthropod body parts that were incorrectly separated during thresholding.

**Analyze Particles**

Using the particle analyzer of ImageJ, this step extracts morphometric measurements (length, width, and area) required for biomass estimation.

Filtering parameters allow exclusion of unwanted particles based on:

- Size (minimum and maximum values interpreted in mm²)
- Circularity (values between 0.0 for a straight line and 1.0 for a perfect circle)

**Crop Image to Petri Dish**

This option restricts the analysis region to the Petri dish containing the specimens.

With the button “Localize Petri Disch” the Petri dish is identified automatically through the following procedure:

1. Generate a binary image using automatic thresholding.
2. Fill holes to remove the dish content.
3. Perform particle analysis.
4. Select the particle with the largest area (assumed to be the Petri dish).
5. Fit a circular region to this particle.
6. Scale and optionally shift the region to exclude the dish edges.

The resulting ellipse defining the Petri dish is specified by width, height, and x/y coordinates. If the crop still does not align with the Petri dish, it is possible to manually change the coordinates and size of the crop region.

**Scales**

To ensure accurate morphometric measurements, all images must be calibrated to convert pixel units into millimeters. This scaling procedure is typically performed using a reference object of known length, such as a metric ruler, included in the same imaging setup as the sample. An image containing the calibration scale is first acquired under the same camera settings and distance as the specimens. The known distance on the ruler (e.g. 10 mm) is then measured in pixels using image analysis software such as ImageJ. From this, a conversion factor (pixels per millimeter) is calculated and applied to all subsequent measurements. This approach ensures that all derived size estimates are standardized and comparable across images and datasets.

**Allometric Regression Models**

Before starting the analysis, users need to define and select the allometric regression parameters for calculating estimations. Users can also:

- Add predefined or new regression models
- Edit existing models
- Remove models from the list

**Finding Fitting Settings**

To display a preview of the processing steps, select one of the loaded images in the pane on the right (Found Images tab). In the newly opened window, the selected image is displayed and on the top a menu bar with different settings is shown. The first drop-down menu (Fig. A4.2) allows viewing the different image processing steps of the pipeline. The second drop-down menu (Fig. A4.3) allows choosing one or more overlays.

| 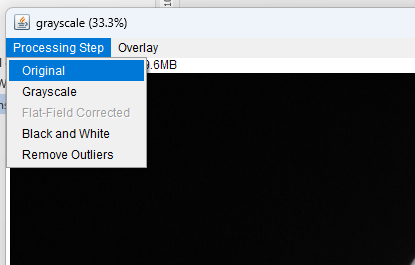  Figure A4.2: Processing steps selectable on single images. | 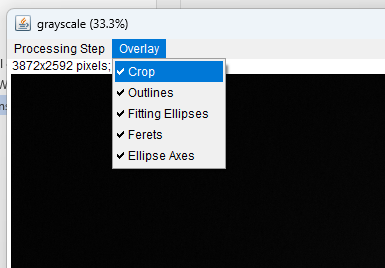  Figure A4.3: Overlay switching in single images. |
| --- | --- |

Following this, the overlays for the Outlines, the Fitting Ellipses, the Ellipse Axes, and the Ferets can be activated and viewed individually in the preview, which is important to verify measurements. By adjusting the settings for the steps “Threshold”, “Remove Outliers” and “Analyse Particles”, the areas included in the results can be changed as desired. Depending on the taxon, different settings might be necessary to discern foreground from background, exclude the wings and legs of the arthropods and include as much of the arthropod body as possible. Once satisfying settings are found, they can be checked for other images by selecting them and opening them in the preview. Furthermore, the settings can be stored in a properties file for future use with the same species.

**AWET Run Procedure (in Folders tab)**

**Start**

Starts the processing pipeline. All images detected in the source folder are analysed sequentially, and the generated results are stored in the destination folder.

**Stop**

Stops the pipeline after the currently processed image has finished.

The analysis will be performed for all found images and for every single image the results will be saved. These include image files for each processing step and zip files containing the ROIs generated by the particle analyzer. Finally, for each image, a CSV file is created which contains all measurements, the converted measurements, and the calculated weights of every particle, once based on the ferets and once based on the fitting ellipses. As values were calibrated on length, width and area of objects. Note that weights calculated based on fitting ellipses are generally more accurate than those for ferets.

**ROI Viewer – Preview of Results (in Roi Viewer tab)**

The ROI Viewer allows users to visualize the regions of interest (ROIs) generated during the particle analysis.

To display ROIs:

1. Select the corresponding ROI archive file (.roi.zip).
2. Select the associated image file.
3. Click View.

A new window will open displaying the image with all detected ROIs labelled and overlaid.

### A5 – Image capture installation

To use the tool on large numbers of images, ensure that the images have the desired region of analysis located in the same place. For this, it is recommended to install a fixed photography station with a Petri dish holder (Fig. A5.1).

Images were acquired using a 10-megapixel digital camera equipped with a standard 50 mm lens and an external flash unit triggered remotely. The flash illuminated the Petri dish from below through a white diffusion box, producing homogeneous lighting conditions. The Petri dish used for imaging had a diameter of 14 cm.

At the end, store the images taken of different taxonomic groups in separate folders, as they may need to be processed separately with different allometric regression models.

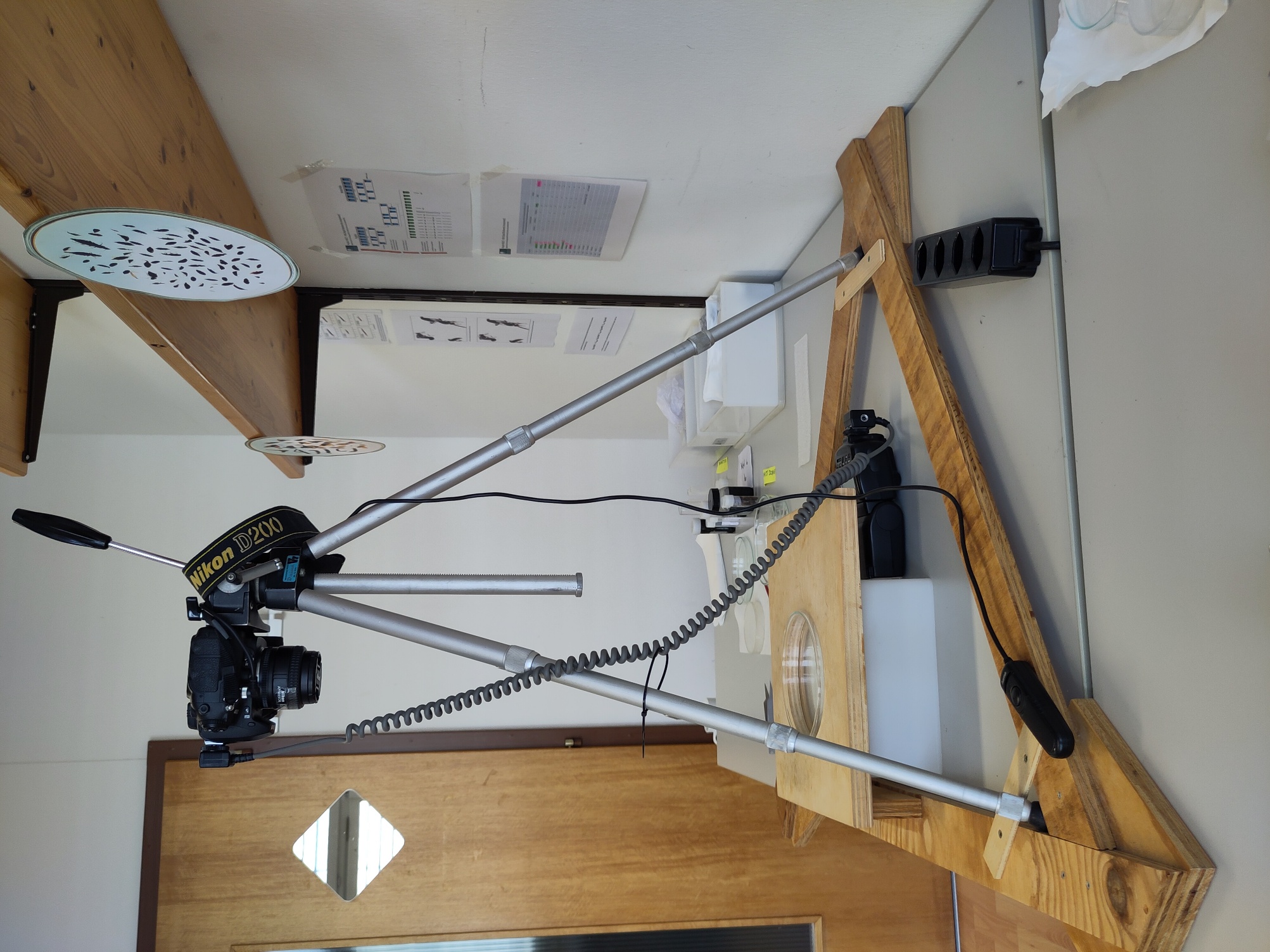

Figure A5.1: Photographic setup used for AWET processing, employing a 10-megapixel digital camera with a standard 50 mm lens and an external flash unit triggered remotely and distributed from below through a diffusion box to the petri dish.
